# Effect of CSFV on Differential Genes of Histone Lactylation at H3K18 in the PI3K-AKT Signaling Pathway

**DOI:** 10.64898/2026.06.26.734696

**Authors:** Huazhou Zhang, Zaiyong Han, Xiaoxia Zhao, Jinhao Zhu, Nan Shao, Kedian Sun, Weiyi Li, Yifei Yao, Xiaowen Liang, Mengru Yang, Yingxue Gao, Jingyi Chen, Yinuo Liang, Xiangdi Li, Qiyuan Liu, Zhi Cao

## Abstract

Classical swine fever (CSF) is a highly contagious disease caused by Classical swine fever virus (CSFV), posing a serious threat to the global swine industry. This study aimed to investigate the effect of CSFV on differential genes of histone lactylation at the H3K18 site in the PI3K-AKT signaling pathway. The site with the most significant change in histone lactylation antibody level was screened by Western blot. Omics analysis was performed using CUT&Tag technology to identify differential genes in the PI3K-AKT pathway between the CSFV-infected group and the mock group, followed by validation using RT-qPCR. Functional analysis of significantly differential proteins was conducted, and the protein expression level of THBS4 was detected by Western blot. The results showed that after CSFV infection of 3D4/21 cells, the H3K18la site exhibited the most significant difference in antibody level. A total of 8,859 differential genes at the H3K18la site were identified by CUT&Tag analysis, including 6,349 up-regulated genes and 2,510 down-regulated genes. Further focusing on the PI3K-AKT signaling pathway, 10 differential genes were identified, comprising 6 up-regulated genes and 4 down-regulated genes. Compared with the control group, the mRNA expression levels of CD19, LAMA1, PDGFRA, BDNF, ANGPT4, and THBS4 were up-regulated in the CSFV-infected group, while FOXO3 and NRTN were down-regulated. Western blot results showed that the protein expression level of THBS4 increased after CSFV infection. These findings lay an important foundation for understanding the molecular mechanisms regulating viral replication and immune evasion, and have significant scientific implications and potential application value.

---

Classical swine fever (CSF) is a highly contagious and often fatal disease that poses a serious threat to pig herds and causes substantial economic losses to the global swine industry. The disease is caused by Classical swine fever virus (CSFV). Although the origin of CSFV remains unclear, many studies suggest that it may be associated with natural infections occurring in boars[1].The virus is primarily transmitted through direct contact, drinking water, and contaminated materials, with clinical manifestations involving various symptoms including respiratory, neurological, and digestive system disorders. CSFV belongs to the genus Pestivirus within the family Flaviviridae, and is a single-stranded positive-sense RNA virus with a genome length of approximately 12.3 kb[2,3].The two ends of the genome are the 5’UTR and 3’UTR untranslated regions, respectively, with a large open reading frame (ORF) in between[4].CSFV encodes multiple proteins, including four structural proteins (C, Erns, E1, and E2) and eight non-structural proteins (Npro, p7, NS2, NS3, NS4A, NS4B, NS5A, and NS5B)[5,6]

During CSFV infection, in addition to the direct interactions between viral proteins and host factors and the impact on a series of host signal transduction pathways, the virus also influences host cell metabolic activities and epigenetic changes. Epigenetic regulation can reprogram host gene expression, thereby facilitating viral replication, immune evasion, and persistent infection. A key feature of viral infection is the remodeling of host cell metabolism. Numerous DNA and RNA viruses can induce the Warburg effect in host cells, a state where cells preferentially undergo glycolysis for energy production even under aerobic conditions, resulting in the abundant production of lactate. This lactate, in turn, promotes the lactylation modification of both histone and non-histone proteins [7,8]. In recent years, with the rise of metabolic-epigenetic research, histone lactylation, as a newly discovered post-translational modification, directly links cellular metabolism (glycolysis and lactate metabolism) with transcriptional regulation, providing a new perspective for exploring how infectious pathogens influence gene expression through metabolic reprogramming [8]. Porcine alveolar macrophages are the primary target cells of CSFV and one of the main effector cells of the immune response. The function of macrophages affects both CSFV proliferation and the host’s immune defense process[9,10].The PI3K/AKT signaling pathway is involved in various physiological activities of macrophages, including activation, energy metabolism, inflammatory responses, and autophagy [11], and plays a critical role during CSFV infection by providing a favorable microenvironment for viral proliferation and transmission.

CSFV NS5A, a multifunctional phosphoprotein, is central to regulating this pathway and can interact with PI3K/AKT signaling through multiple mechanisms. Specifically, NS5A binds to the autophagy-related protein Beclin1 and upregulates its expression, thereby modulating the activity of the PI3K/AKT/mTOR signaling axis to promote CSFV replication[12]. However, the precise molecular mechanisms by which CSFV regulates the PI3K/AKT pathway and histone lactylation through metabolic and epigenetic reprogramming remain unclear.

Histone lysine lactylation occurs at multiple sites; among them, lysine 18 of histone H3 (H3K18) is located in the histone tail and is closely associated with transcriptional activation[13]. H3K18 lactylation influences the activation of transcription factors and RNA polymerase in promoter regions by altering chromatin conformation, thereby regulating the expression levels of genes involved in downstream signaling pathways [14]. During viral infection, infection-induced enhancement of glucose metabolism and increased lactate production provide abundant metabolic substrates for H3K18 lactylation; these metabolic changes may affect the transcriptional regulation of PI3K/AKT pathway genes via lactylation-mediated epigenetic modifications[15,16].

In this study, we utilized CUT&Tag—a simple, reproducible, and highly sensitive technology—to screen for differentially enriched genes at the H3K18la site, with particular emphasis on genes involved in the PI3K-AKT signaling pathway. Combined with RT-qPCR validation, we explored the effects of CSFV infection on the expression of PI3K-AKT pathway-related genes in porcine macrophages. These findings provide a foundation for further elucidating the molecular mechanisms underlying viral replication and immune evasion.

## 2. Materials and methods

### 2.1 Major Reagents

DL5000 DNA Marker, PAGE Gel Fast Preparation Kit, Fetal Bovine Serum (FBS), RNA Extraction Kit, 2×AceQ qPCR SYBR Green Master Mix, and Reverse Transcriptase were purchased from Nanjing Vazyme Biotech Co., Ltd. Goat Anti-Rabbit IgG H&L pAb (rabbit secondary antibody), Anti-L-Lactyl-Histone H3 (H3K18) Rabbit mAb-ChIP Grade, L-Lactyl Lysine, H3K9la, H3K14la, H2B, H3K18la, H4K8la, H4K12la, H2A, H3K23la, H4K16la, Histone 3, and β-actin antibodies were purchased from Hangzhou Jingjie Biotechnology Co., Ltd. Ultrapure water (DNase and RNase-free), HRP-Goat Anti-Rabbit IgG, and HRP-Goat Anti-Mouse IgG were purchased from Beyotime Biotechnology (Shanghai) Co., Ltd. The sequencing kit NovaSeq X Series 25B Reagent Kit (300 Cycle) was purchased from Illumina, Inc. The BCA Protein Assay Kit, non-fat dry milk, 5 × SDS Protein Loading Buffer, and Femto-sig ECL Chemiluminescence Substrate Kit were purchased from Shanghai Epizyme Biomedical Technology Co., Ltd. The NovoNGS® CUT&Tag High-Sensitivity Kit and NovoNGS® Cut&Tag Illumina Primer Kit were purchased from Suzhou Novoprotein Scientific Co., Ltd.

### 2.2 Cells

The NS3-NS4A plasmid, NS3 monoclonal antibody, CSFV viral stock, and porcine alveolar macrophages were all preserved in our laboratory. RPMI 1640 medium and 1% penicillin-streptomycin (dual antibiotics) were purchased from HyClone. 3D4/21 cells were cultured in RPMI 1640 medium supplemented with 10% fetal bovine serum (purchased from Vazyme Biotech Co., Ltd.) and 1% dual antibiotics at 37 °C in a 5% CO₂ atmosphere.

### 2.3 Western Blot Detection of Histone Antibody Levels

Cells collected at 24 h, 48 h, 60 h, and 72 h post-CSFV infection were scraped using a cell scraper, resuspended in 1 mL of ice-cold PBS, and transferred to 1.5 mL centrifuge tubes. After centrifugation (4 °C, 12,000 rpm, 10 min), the cell pellets were collected and washed four times. Histones were extracted using a histone extraction kit. After BSA protein quantification, histone antibody levels were detected by Western blot.

### 2.4 Screening of Differential Genes in the PI3K-AKT Pathway by CUT&Tag

Cell samples from the CSFV-infected group and the blank control group were collected at 48 h post-infection using a cell scraper. After resuspension in ice-cold PBS, cells were centrifuged at 4 °C and 12,000 rpm for 10 min, and the washing step was repeated four times. The cell pellets were then gently washed with an equal volume of 1×Wash Buffer and centrifuged. The cell pellets were resuspended in 90 μL of 1×Wash Buffer. ConA magnetic beads were washed with 1× ConA Binding Buffer, and 10 μL of the washed beads were added to the cell suspension, followed by incubation with rotation at room temperature for 10 min. After magnetic separation, 50 μL of ice-cold Primary Antibody Buffer containing 1 μg of primary antibody was added, and the mixture was incubated statically at 4 °C overnight. Subsequently, 90 μL of secondary antibody diluted 1:200 was added, and the mixture was incubated with rotation at room temperature for 1h. The beads were washed twice with Antibody Buffer. Then, 72 μL of transposome dilution buffer containing ChiTag® pAG-Transposome was added, followed by incubation with rotation at room temperature for 1h. After washing, 65 μL of Tagmentation Buffer containing 0.7 M MgCl₂ was added, and the reaction was incubated at 37 °C for 1h to perform tagmentation. The reaction was terminated by adding 5 μL of Stop Buffer and incubating at 55 °C for 10 min. Subsequently, 144 μL of Tagment DNA Extract Beads were added for DNA extraction. After ethanol washing and elution with Elution Buffer, 1 μL of the eluted DNA was used for pre-amplification (with cycle numbers of 17, 20, and 23 cycles, respectively) followed by nucleic acid gel electrophoresis. The PCR products were purified, and the purified products were subjected to sequencing. The obtained sequencing data were analyzed using bioinformatics methods.As shown in Table 1.

**Table 1.** PCR Amplification system.

| Component | Volume |
| --- | --- |
| DNA template | 20.0 μL |
| i5 Primer(10 μM) | 2.5 μL |
| I7 Primer(10 μM) | 2.5 μL |
| 2×HiFi AmpliMix | 25.0 μL |
| Total | 50.0 μL |

### 2.5 Quantitative Real-Time PCR (RT-qPCR) Analysis of Differential Gene Expression at the mRNA Level

Cells were harvested at 48 h and 60 h following CSFV infection or transfection with the NS3-NS4A plasmid. Total RNA was extracted, and reverse transcription was performed to obtain cDNA. The mRNA expression levels of target genes were quantified by qRT-PCR using the following reaction mixture 20 μL: 10 μL of 2×AceQ qPCR SYBR Green Master Mix, 1 μL of cDNA, 0.5 μL of forward primer, 0.5 μL of reverse primer, and 8 μL of nuclease-free water. The amplification program consisted of an initial denaturation step at 95 °C for 5 min, followed by 40 cycles of 95 °C for 10 s and 60 °C for 30 s.As shown in Table 2.

**Table 2.**
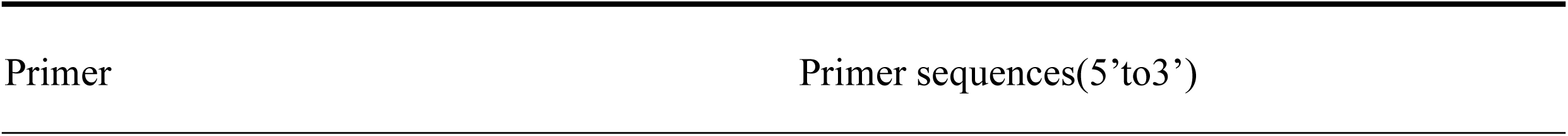
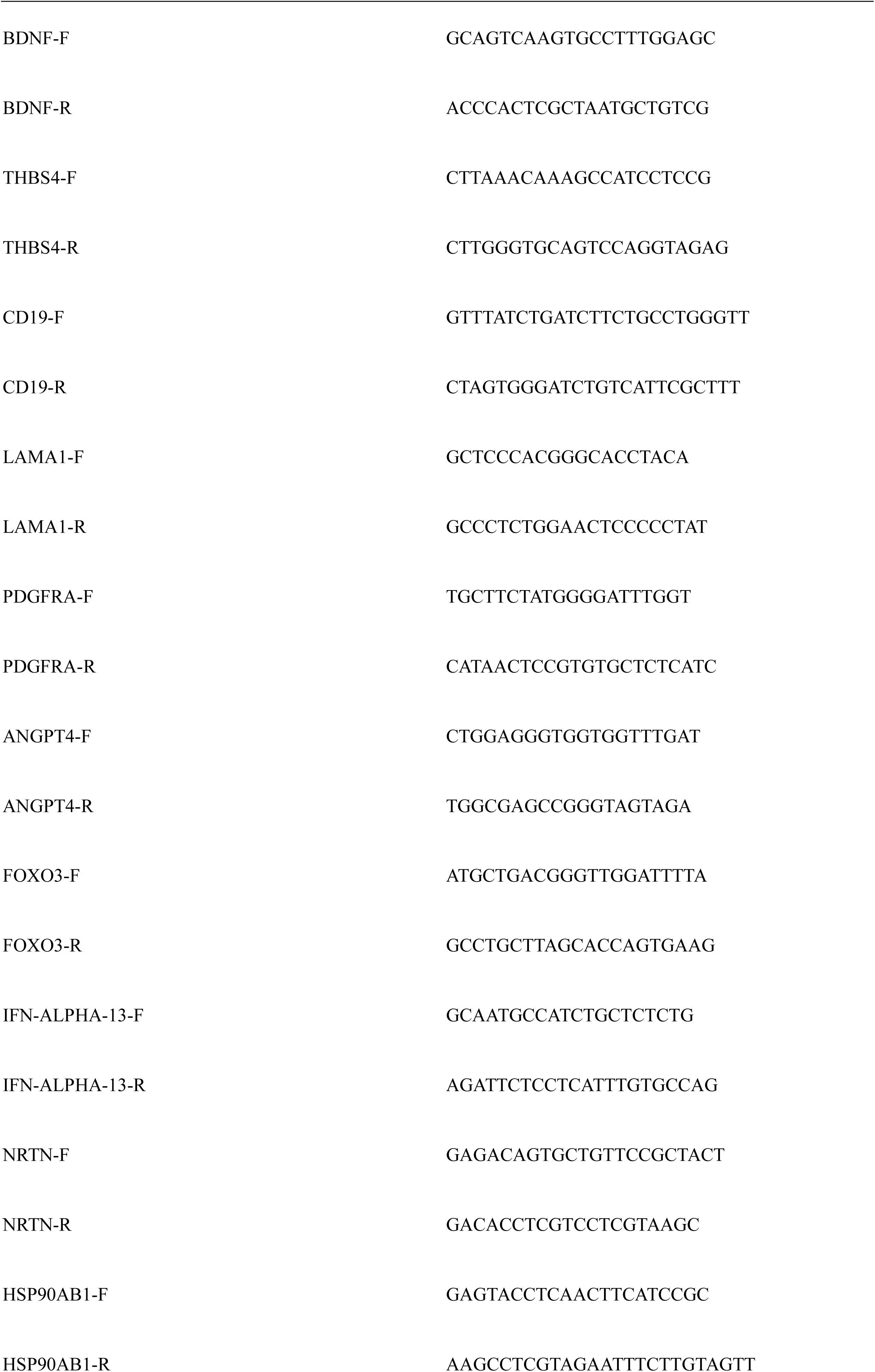
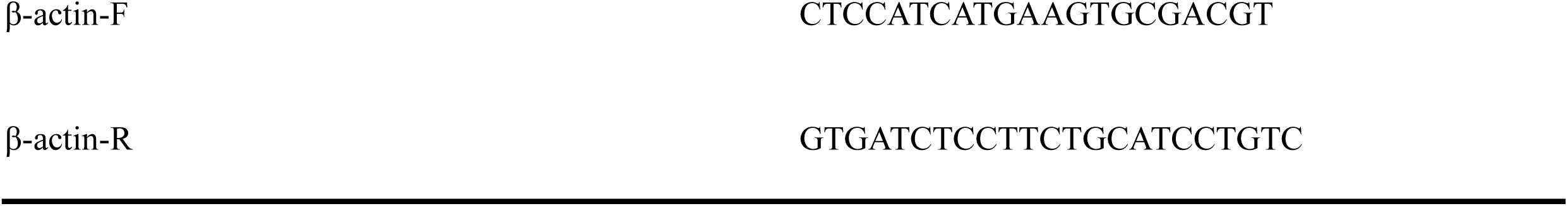
Primer sequences of differentially expressed genes.

### 2.6 Western Blot Analysis of Key Protein Expression

3D4/21 cells were trypsinized, resuspended, and adjusted to a density of 1.0 × 10⁶ cells/mL before seeding into culture plates. Upon reaching approximately 80% confluence, the medium was replaced with low-serum medium (2% FBS). Cells were then infected with CSFV and incubated in a CO₂ incubator. At 48 h post-infection, cells were collected and lysed with RIPA lysis buffer on ice for 30 min, followed by centrifugation at 12 000 rpm for 10 min. The supernatants were harvested, and protein samples were analyzed by Western blotting to assess the expression levels of the key proteins.

## 3. Results

### 3.1 Analysis of Cellular Pan-Antibody Lactylation Levels by Western Blot

Following CSFV infection of porcine alveolar macrophages, cell samples were harvested at 12h, 24h, 48h, and 60h post-infection. Histones were extracted, and subsequent Western blot analysis using a pan-lactylation antibody showed a significant increase in lactylation levels.As shown in Figure 1.

**Fig. 1.**
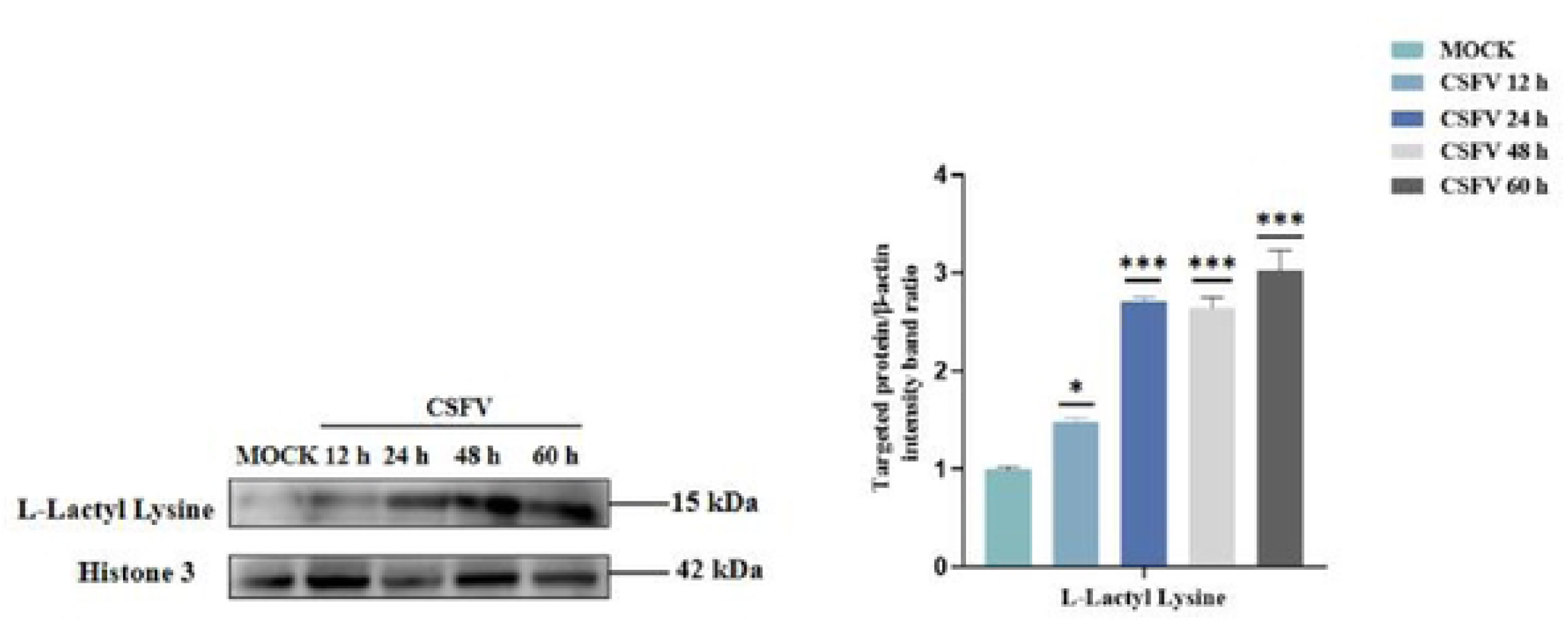
Global histone lactylation levels in CSFV-infected cells.Histones were extracted from CSFV-infected 3D4/21 cells at different time points and analyzed using a pan-lactylation antibody by Western blot. CSFV infection increased overall histone lactylation levels compared with the mock group.

### 3.2 CSFV Infection Enhances Cellular Lactylation Antibody Levels

The protein levels of H3K9la, H3K14la, H2B, H3K18la, H4K8la, H4K12la, H2A, H3K23la, H4K16la, and total Histone 3 were further examined by Western blot. CSFV infection was found to increase cellular lactylation levels. Among these, H3K18la showed the greatest difference and was therefore selected for CUT&Tag analysis of histone modifications.As shown in Figure 2.

**Fig. 2.**
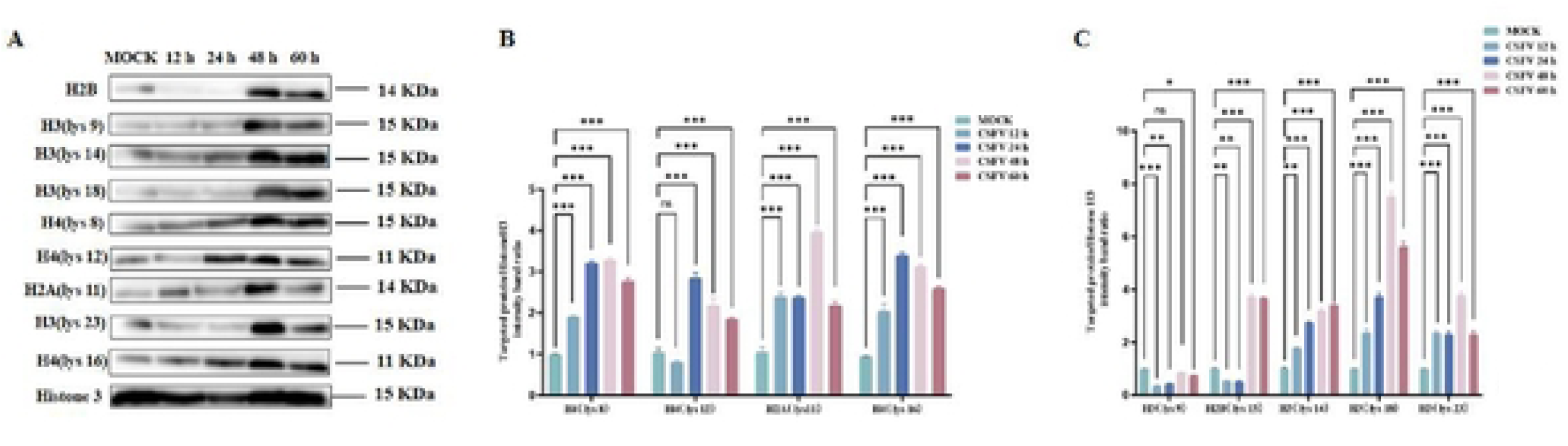
Detection of histone lactylation modification sites following CSFV infection.The levels of multiple histone lactylation sites were analyzed by Western blot. H3K18la exhibited the most significant increase and was selected for subsequent CUT&Tag analysis.

### 3.4 Sample Analysis Results

Sample reproducibility within groups was assessed by calculating the correlation of sequencing read distribution across the reference genome. The reference genome was binned into 10 kb intervals, and the number of reads mapped to each interval was counted per sample. Following log2 transformation, correlation coefficients between samples were computed and visualized as a heatmap. The results demonstrated good reproducibility and high correlation among samples. Sequencing reads were predominantly centered at peak regions, with clear signal enrichment, indicating high data quality.As shown in Figure 3.

**Fig. 3.**
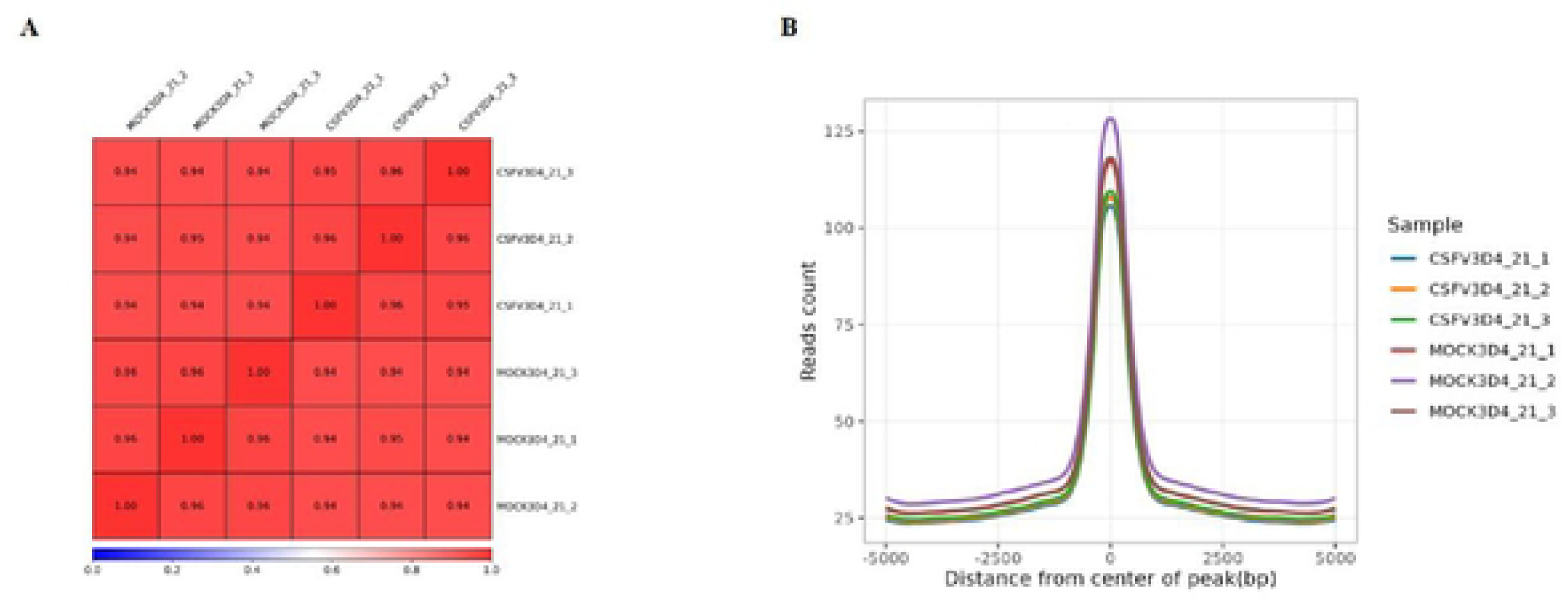
Quality assessment and reproducibility analysis of CUT&Tag sequencing data.(A) Correlation heatmap showing Pearson correlation coefficients among biological replicates from the CSFV-infected and mock groups. The reference genome was divided into 10 kb bins, and sequencing read counts within each bin were calculated and log2-transformed before correlation analysis.(B) Signal enrichment profiles centered around peak regions identified by CUT&Tag sequencing. Sequencing reads displayed significant enrichment near peak centers with low background noise, indicating high-quality library construction and reliable sequencing performance.The results demonstrated strong reproducibility among biological replicates and confirmed the suitability of the sequencing data for downstream differential peak analysis.

### 3.5 Statistical Results of Differential Peaks

To explore differences in the binding of transcription factors or histone modifications to DNA under distinct conditions and to elucidate the regulatory functions and dynamic mechanisms of epigenetic regulation, the Peak position files from different sample groups were merged using bedtools. The read counts within the merged regions were calculated for each sample, followed by differential analysis to obtain differential Peaks between sample groups. The filtered differential Peaks were subsequently subjected to downstream analyses, including motif prediction, Peak annotation, and functional enrichment analysis. In the comparison between the CSFV-infected group and the MOCK group, 6 349 up-regulated Peaks and 2 510 down-regulated Peaks were identified.As shown in Figure 4.

**Fig. 4.**
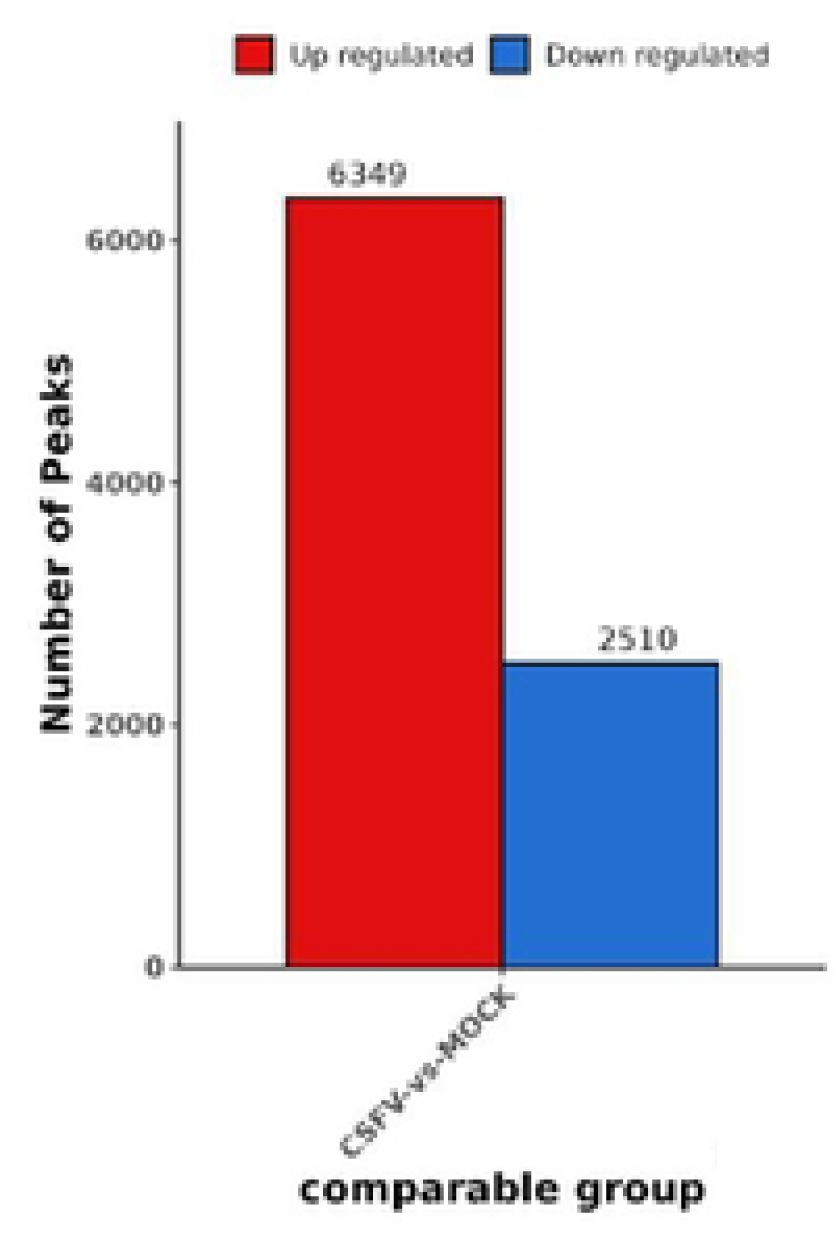
Differential peak analysis between CSFV-infected and mock groups.A total of 8,859 differential peaks were identified, including 6,349 up-regulated peaks and 2,510 down-regulated peaks.

#### 3.5.1 Functional Region Annotation of Differential Peaks

The binding of target proteins to different genomic regions can regulate gene expression through distinct mechanisms. Functional annotation of Peaks facilitates the analysis of the regulatory mechanisms of target proteins. In this study, the R package chipSeeker was used to annotate Peaks from each sample. The distribution of all Peaks across various functional regions was statistically analyzed and visualized as a bar chart in the following order: Promoter > 5’UTR > 3’UTR > Exon > Intron > Downstream > Intergenic. The results showed a similar overall distribution pattern across samples, indicating stable data quality.As shown in Figure 5.

**Fig 5.**
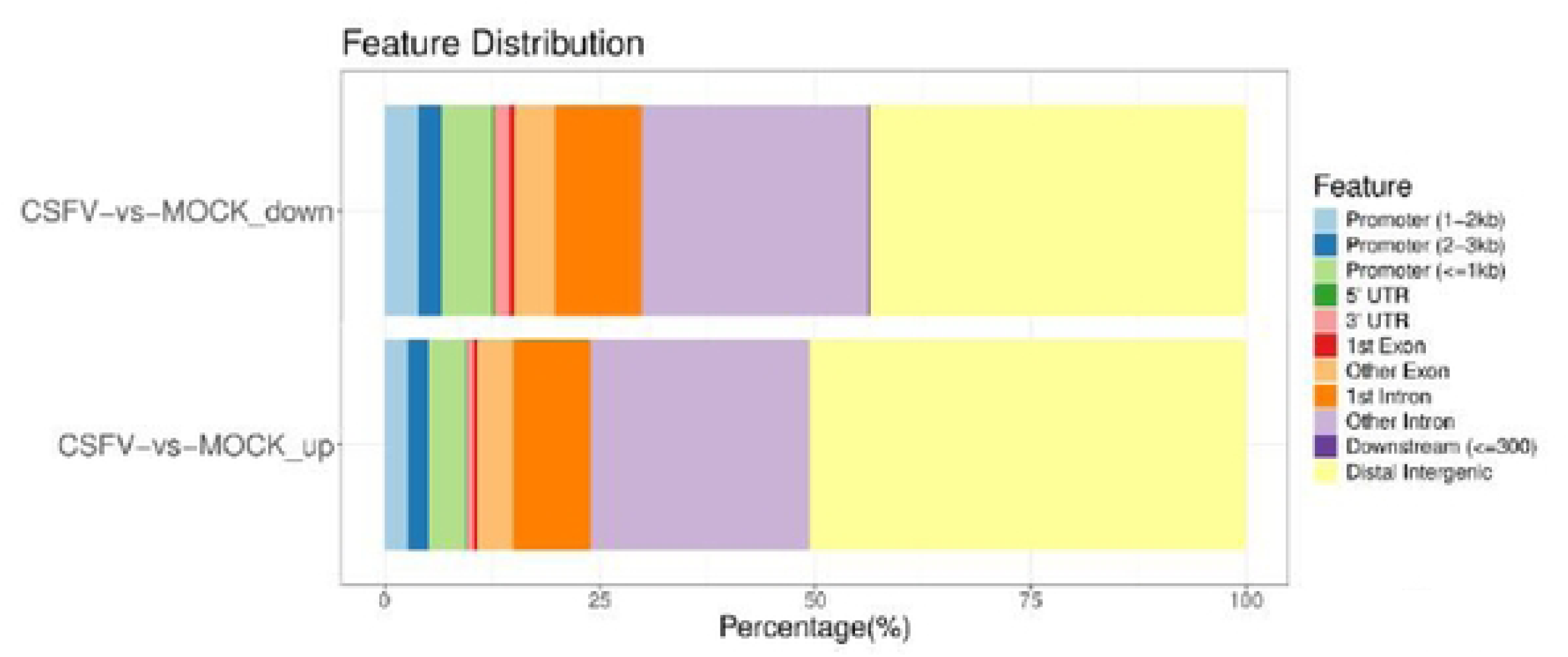
Functional annotation of differential peaks.Differential peaks were annotated according to genomic regions. Most peaks were distributed within promoter regions.

#### 3.5.2 GO Functional Enrichment Analysis of Genes Neighboring Differential Peaks

GO enrichment analysis of genes located near differential Peaks allows the identification of significantly enriched GO terms, thereby revealing the functional differences in target protein regulation between the two groups. GO annotation was performed for genes adjacent to each differential Peak. The significance levels of gene enrichment for each GO term were calculated across three ontologies: Biological Process (BP), Cellular Component (CC), and Molecular Function (MF). GO terms with a q-value < 1 were retained for subsequent analysis. The analysis revealed that the enriched GO functions primarily involved biological processes, cellular components, and molecular functions.As shown in Figure 6.

**Fig 6.**
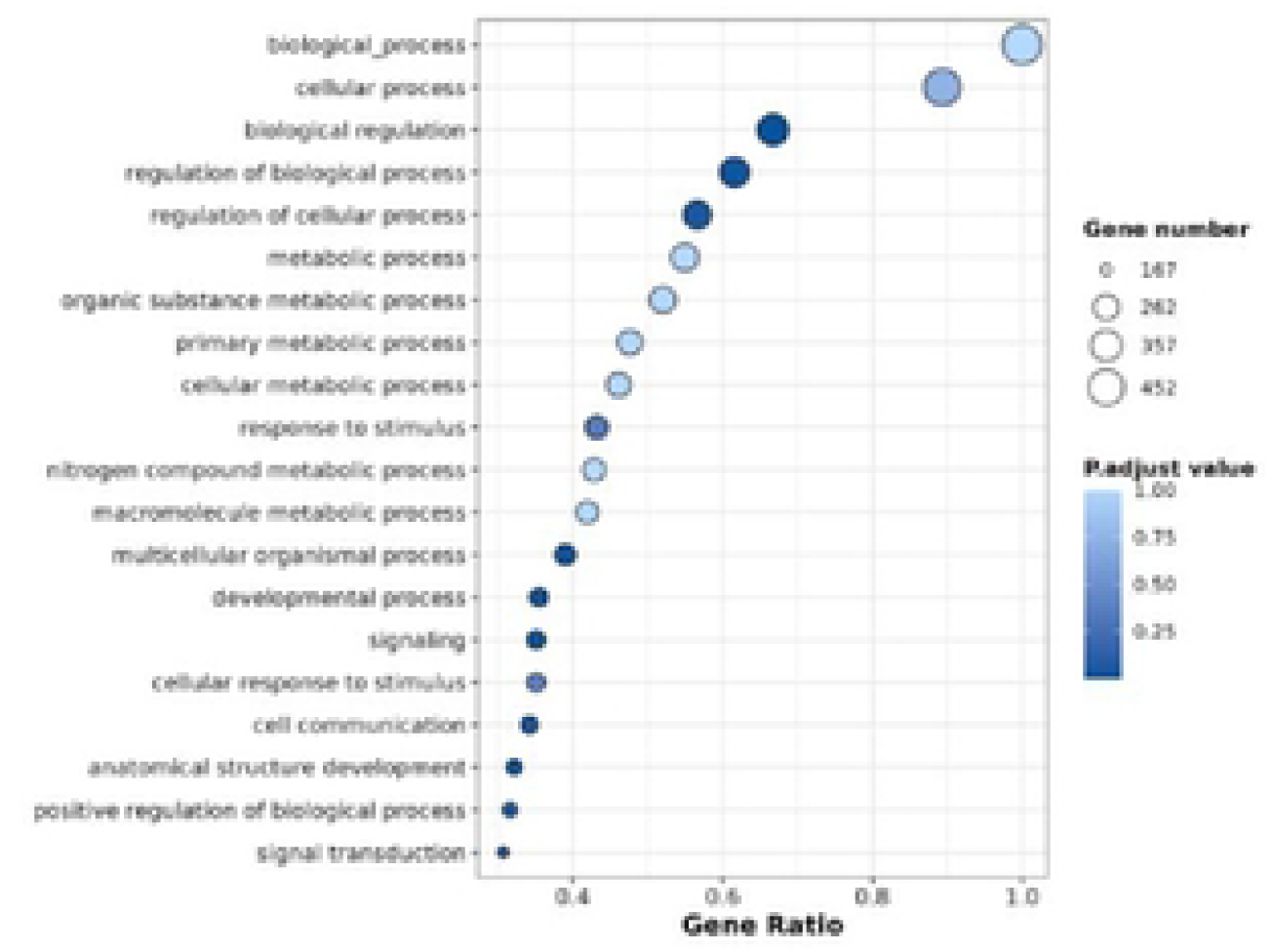
GO enrichment analysis of genes associated with differential peaks.Genes neighboring differential peaks were subjected to GO enrichment analysis. The enriched terms were mainly related to biological processes, cellular components, and molecular functions.

#### 3.5.3 KEGG Enrichment Analysis of Genes Neighboring Differential Peaks

KEGG pathway enrichment analysis of genes located near differential Peaks allows the identification of significantly enriched KEGG pathways, thereby revealing the metabolic pathways that are differentially regulated by the target protein between the two groups. To this end, KEGG functional annotation was first performed for genes adjacent to each differential Peak. Subsequently, the significance level of gene enrichment for each KEGG pathway was calculated, and pathways with a q-value < 1 were selected for further analysis. The results demonstrated that the PI3K-AKT pathway exhibited a relatively large number of neighboring genes and showed significant pathway enrichment.As shown in Figure 7.

**Fig. 7.**
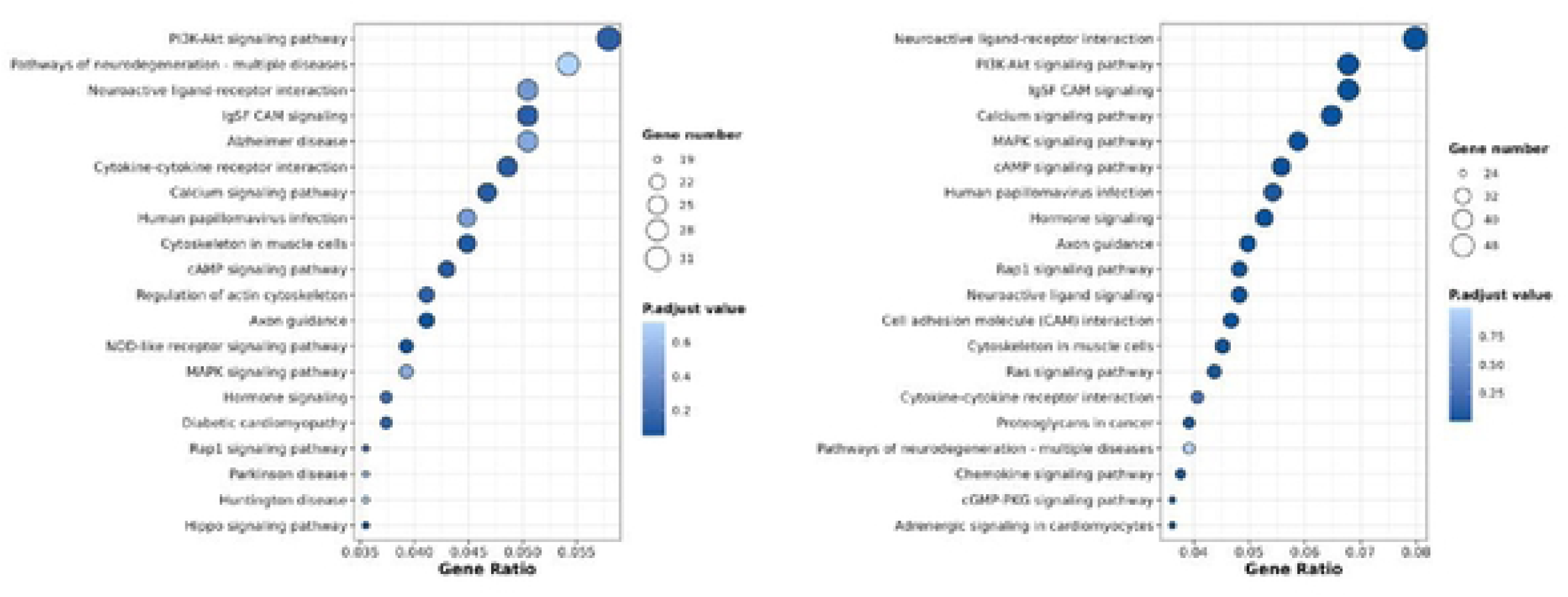
KEGG pathway enrichment analysis of genes associated with differential peaks.KEGG analysis identified multiple significantly enriched pathways. The PI3K-AKT signaling pathway was selected for further investigation.

#### 3.5.4 Analysis of Differential Genes in the PI3K-AKT Pathway

Further screening targeting the PI3K-AKT signaling pathway and promoter regions revealed 10 differentially expressed genes between the CSFV experimental group and the control group (Table 3), including 6 up-regulated genes and 4 down-regulated genes.

**Table 3.**
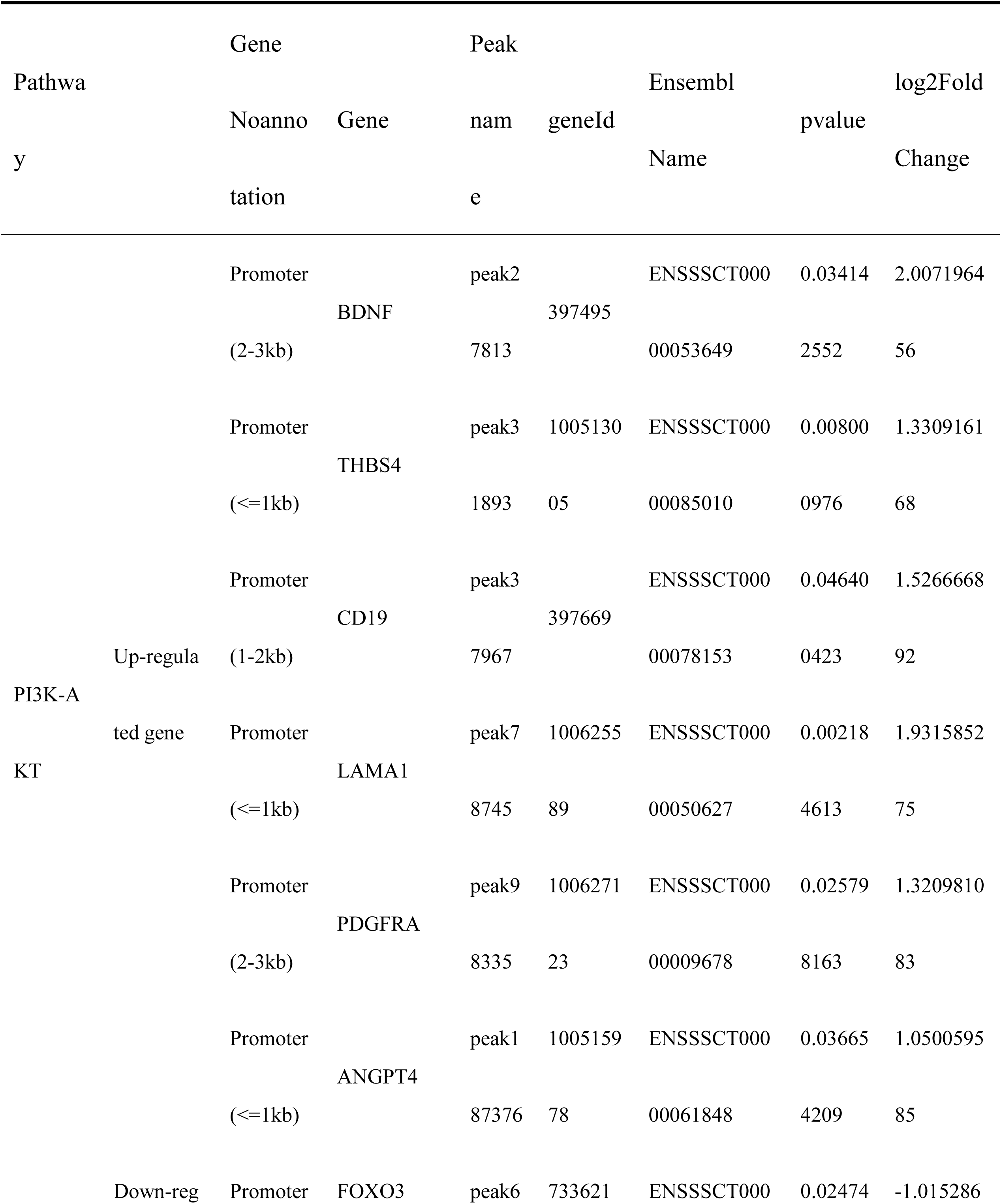
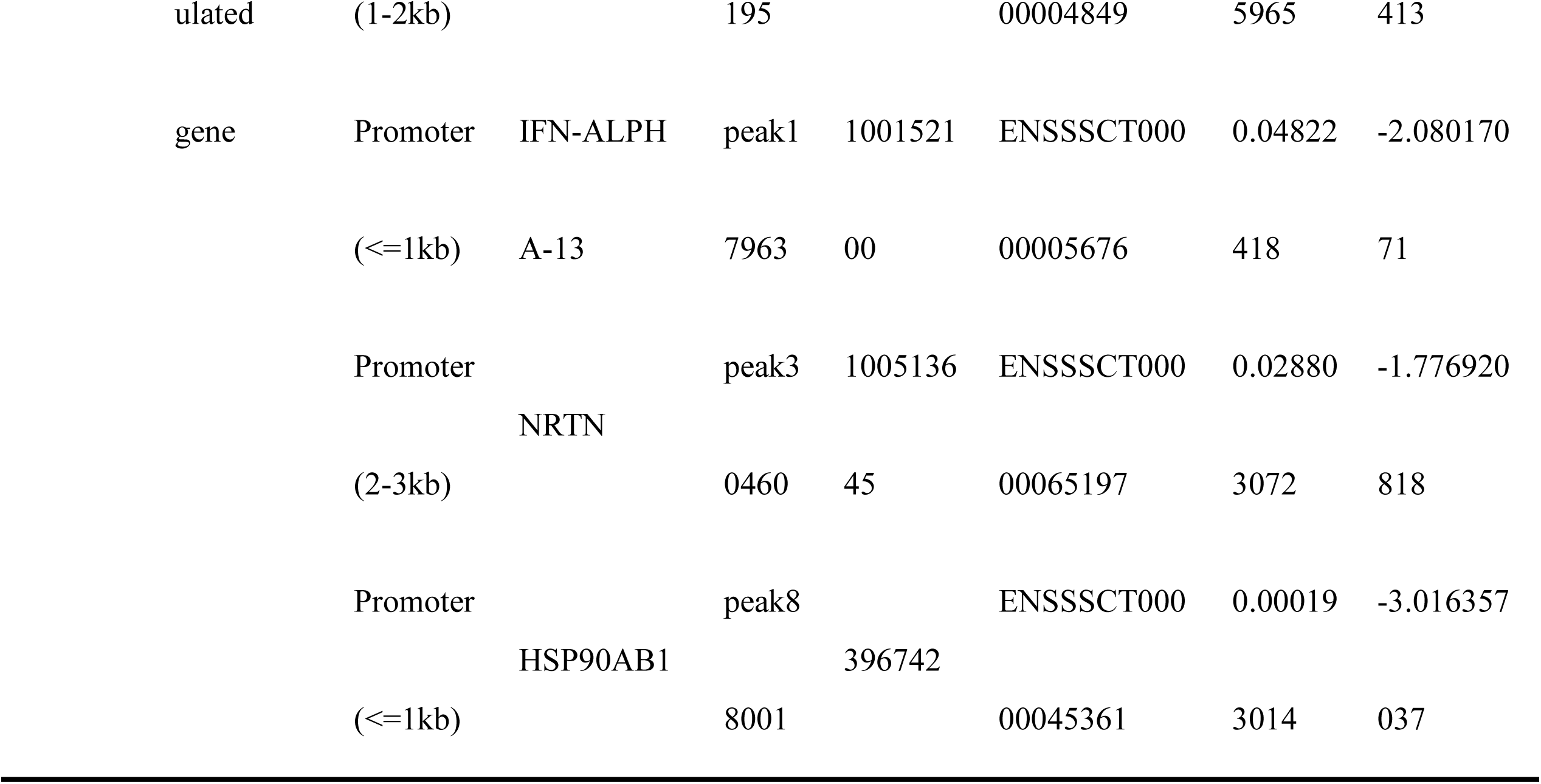
Differential genes in the PI3K-AKT signaling pathway.

### 3.6 CSFV Alters the Expression of Differential Genes in the PI3K-AKT Pathway in Porcine Alveolar Macrophages

To determine the effect of CSFV on the expression of differential genes involved in the PI3K-AKT pathway, 3D4/21 cells were infected with CSFV, and samples were collected at 48 h and 60 h post-infection. The expression levels of the target genes were validated by qPCR. The results showed that the expression levels of CD19, LAMA1, PDGFRA, BDNF, ANGPT4, and THBS4 were up-regulated, whereas FOXO3 and NRTN were down-regulated.As shown in Fig 8.

**Fig. 8.**
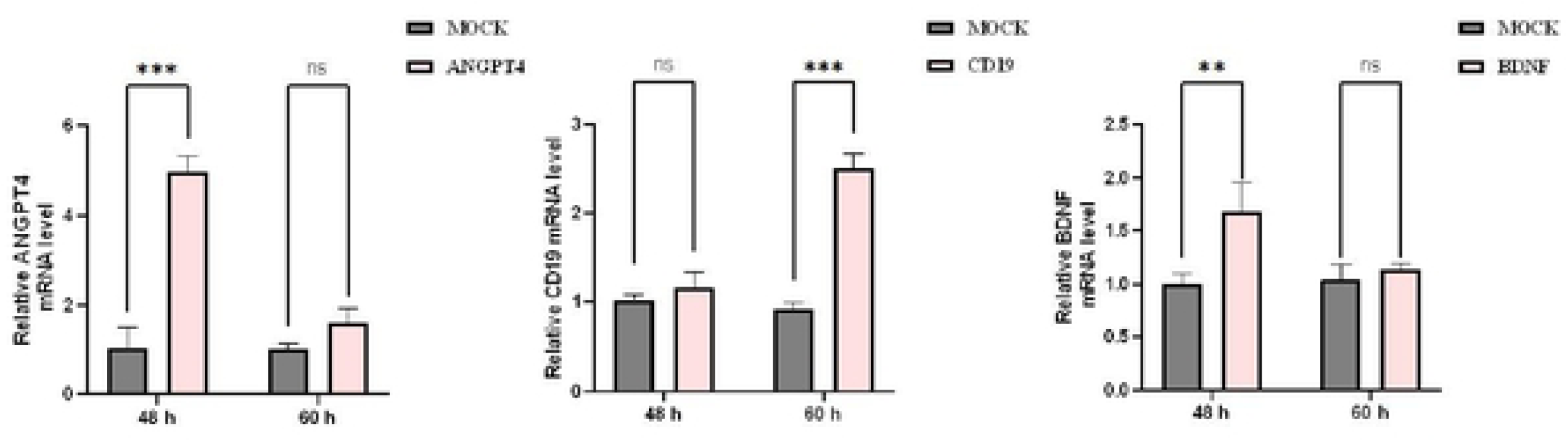
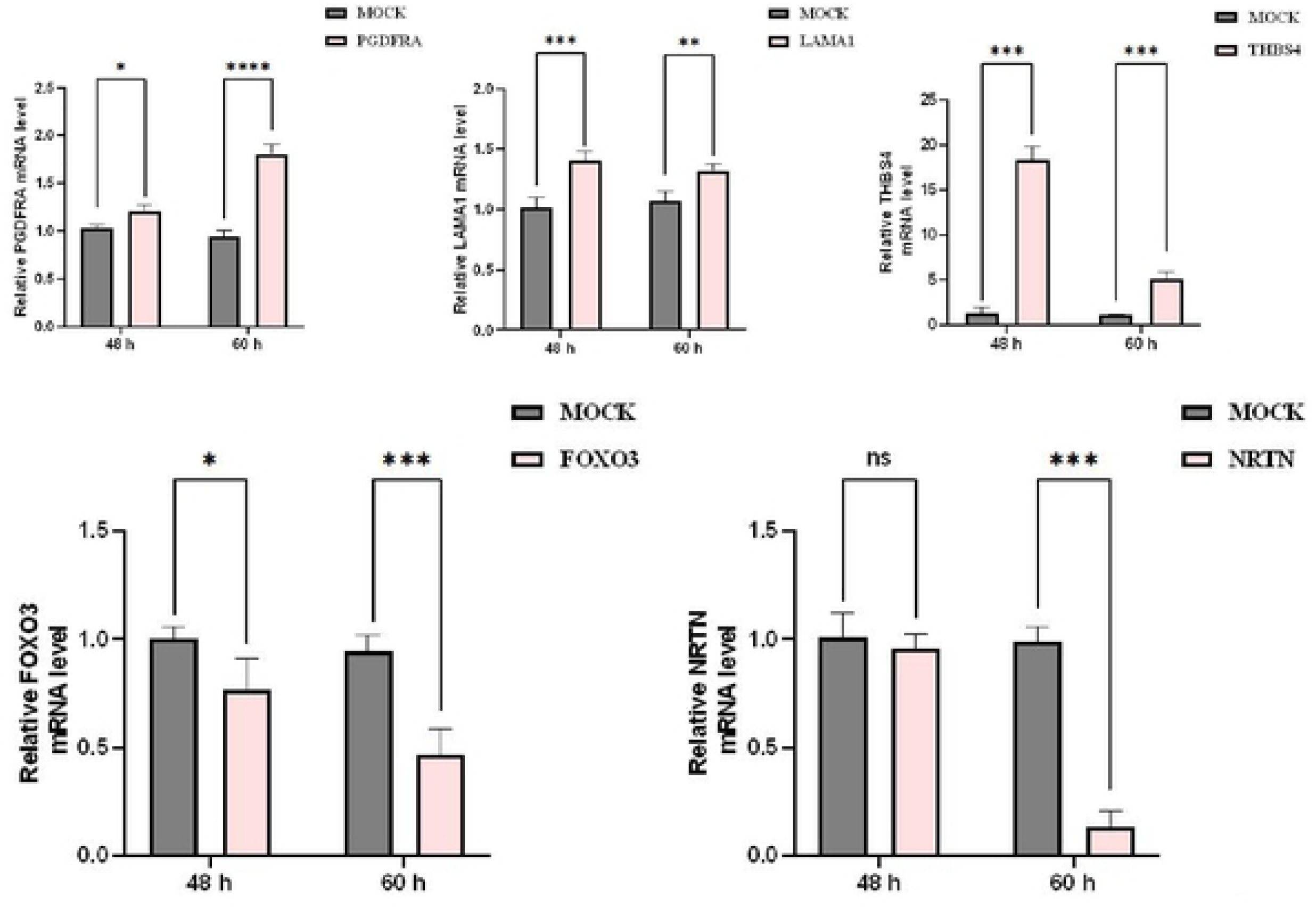
qPCR analysis of differential gene expression in the PI3K-AKT signaling pathway upon CSFV infection Note: Data are shown as the mean values ± SD of three independent experiments.*\* p <*0.05, ** *p* <0.01, *** *p* <0.001, were calculated using the t-test for NS3-NS4A infected versus empty (MOCK) infected group.

### 3.7 Western Blot Detection of THBS4 Expression Levels in CSFV-Infected Cells

Based on the effect of CSFV NS3-NS4A on differential genes in the PI3K-AKT pathway, THBS4, which exhibited the most significant difference at the mRNA level, was selected for further investigation. Porcine alveolar macrophages were infected with CSFV, and protein samples were collected at 48 h post-infection for Western blot analysis to detect the endogenous expression level of THBS4. The results showed that the protein expression level of THBS4 was elevated in the CSFV-infected group compared with the mock group.As shown in Figure 9.

**Fig. 9.**
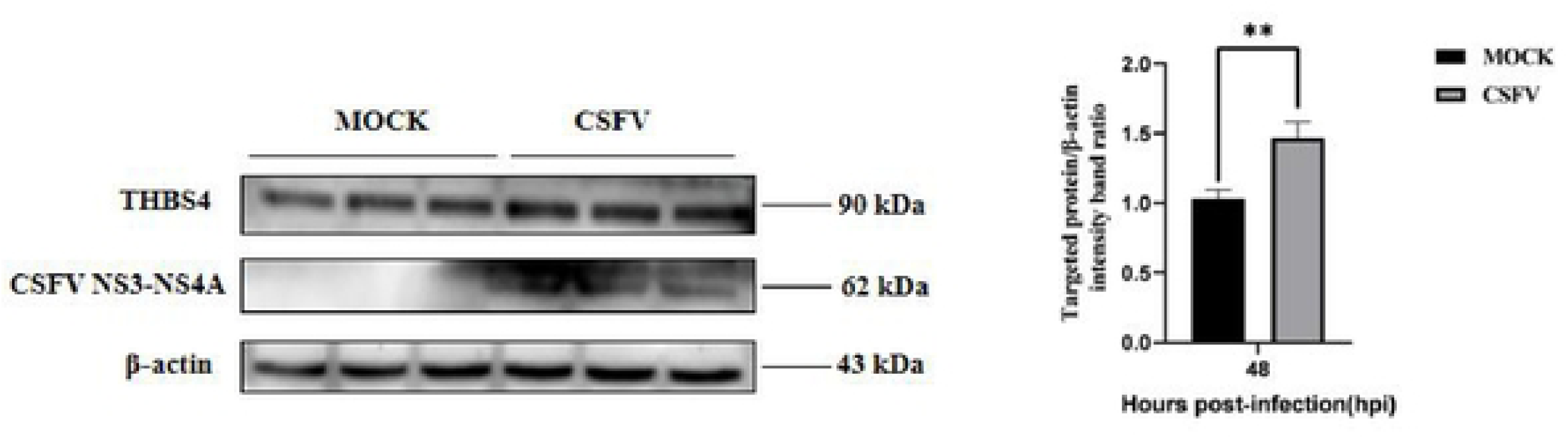
Western blot validation of THBS4 protein expression in CSFV-infected cells.THBS4 was selected as a representative target gene for protein-level validation due to its significant differential enrichment and transcriptional up-regulation following CSFV infection. Total cellular proteins were extracted from CSFV-infected and mock-treated 3D4/21 cells at 48 h post-infection. THBS4 protein expression was detected by Western blot analysis, and β-actin served as the loading control. Densitometric analysis demonstrated a significant increase in THBS4 protein abundance in the CSFV-infected group, consistent with the RT-qPCR and CUT&Tag results. These findings suggest that H3K18la-mediated epigenetic regulation may contribute to THBS4 activation during CSFV infection.Data are presented as mean ± SD from three independent biological replicates.\**p* < 0.05 \*\**p* < 0.01 (Student’s t-test).

## 4 Discussion

CSFV utilizes macrophages as host immune carriers. During CSFV infection, a series of changes occur in host cell signaling pathways, metabolism, and gene expression. In this study, CUT&Tag technology combined with RT-qPCR validation was employed to thoroughly investigate the effect of CSFV on differential genes associated with the histone lactylation H3K18 site within the PI3K-AKT signaling pathway.

CUT&Tag, as an emerging chromatin profiling method, is widely used to precisely map histone modifications and transcription factor binding sites due to its advantages, including ease of operation, high signal-to-noise ratio, and low input requirements[17].These powerful advantages have brought new opportunities to the life sciences field and can be utilized to investigate epigenetic modifications, transcription factor binding sites, chromatin three-dimensional structure, and other aspects[18,19]. In the field of plant science, researchers successfully constructed a cDNA library from coronatine-induced cambial cells of Hevea brasiliensis using CUT&Tag technology. They found that histone H3 acetylation is associated with the enrichment of genes involved in auxin, flavonoid metabolism, and protein ubiquitination, regulating the expression of these genes during secondary laticifer differentiation. This finding provides a theoretical basis for further elucidating the molecular mechanisms by which histone modifications regulate secondary laticifer differentiation in rubber trees[20]; In the field of oncology, a research team used CUT&Tag technology and found that H4K12 lactylation in triple-negative breast cancer (TNBC) binds to the promoter region of SLFN5 and suppresses its expression, thereby promoting malignant tumor progression[21];In the field of parasitology, CUT&Tag technology was applied for the first time to the epigenomic study of the AT-rich genome of Plasmodium falciparum, successfully overcoming the limitations of conventional methods. With as few as 10 000 nuclei, genome-wide maps of heterochromatin (H3K9me3, HP1) were obtained, showing high concordance with ChIP-seq data and extremely low background signals. This breakthrough provides a reliable technical solution for chromatin profiling of low-input samples, and is particularly suitable for parasite stages that are difficult to obtain, such as the mosquito stage and liver stage of Plasmodium[22].These studies fully demonstrate that CUT&Tag technology holds significant research value and broad application prospects across various fields.

In this study, CUT&Tag technology was successfully employed to screen for genes showing significant differential enrichment at the H3K18la site in CSFV-infected cells, with a particular focus on genes associated with the PI3K/AKT signaling pathway. This provides new insights into the role of epigenetic modifications in the pathogenesis of CSFV. The results showed that CSFV infection significantly upregulated the overall level of cellular histone lactylation, with the H3K18la site exhibiting the most pronounced differential enrichment. Previous studies have demonstrated that CSFV infection can upregulate the lactylation level of histone H2B (H2BK16la) in PK-15 cells via the LDHA-lactate axis, thereby regulating the activation of the NF-κB pathway and the expression of type III interferons[23]. This study extends this line of research to the H3K18la site, laying a foundation for the study of lactylation in CSFV. In the differential expression analysis of genes associated with the PI3K-AKT signaling pathway, six genes—CD19, LAMA1, PDGFRA, BDNF, ANGPT4, and THBS4—were found to be significantly up-regulated, whereas four genes—FOXO3, NRTN, IFN-ALPHA-13, and HSP90AB1—were significantly down-regulated. Changes in H3K18la enrichment following CSFV infection correlated with the altered expression of certain PI3K-AKT pathway genes, suggesting that these genes may be direct or indirect targets of H3K18la modification. For instance, among the up-regulated genes, PDGFRA and BDNF are both upstream activators of AKT signaling; their increased expression may enhance pathway activity through a positive feedback mechanism. Among the down-regulated genes, FOXO3, a key downstream transcription factor of AKT, is repressed, which may favor cell survival and anti-apoptosis. The differential genes identified in the PI3K-AKT pathway in this study provide an experimental basis for subsequent proteomic analysis of PI3K-AKT differential proteins regulated by CSFV NS3-NS4A.

## 5. Conclusion

This study demonstrates that CSFV infection upregulates cellular histone lactylation levels, with the most significant difference observed at the H3K18la site. Further analysis using CUT&Tag technology revealed that six genes in the PI3K-AKT pathway were up-regulated and four were down-regulated. Moreover, the protein expression level of THBS4 was significantly increased in CSFV-infected porcine alveolar macrophages. These findings lay a foundation for elucidating the molecular mechanisms regulating viral replication and immune evasion.

## Author Contributions

Conceptualization: HZ Zhang, ZY Han, Z Cao. Data curation: XX Zhao, JH Zhu, N Shao. Formal analysis: KD Sun, WY Li, YF Yao. Investigation: XW Liang, MR Yang, YX Gao. Methodology: HZ Zhang, ZY Han. Project administration: Z Cao. Resources: JY Chen, YN Liang, XD Li, QY Liu. Supervision: Z Cao. Validation: XX Zhao, JH Zhu, N Shao. Visualization: HZ Zhang, ZY Han. Writing – original draft: HZ Zhang, ZY Han. Writing – review & editing: Z Cao, HZ Zhang, ZY Han.

## Availability of data and materials

The datasets supporting the conclusions of this article are included within the article.

## Ethics declaration

All experiments received approval and were supervised by the Research Ethics Committee of Qingdao Agricultural University. Informed consent was obtained from all owners for fecal sample collection. All methods were performed in accordance with the relevant guidelines and regulations. The study did not involve human participants or clinical trials; therefore, no further consent for participation or publication was required.

## References

1. Moennig V. The control of classical swine fever in wild boar. Frontiers in Microbiology, 2015, 6: 1211.

2. Ganges L, Crooke H R, Bohórquez J A, et al. Classical swine fever virus: the past, present and future. Virus Research, 2020, 289: 198151.

3. Fan J, Liao Y, Zhang M, et al. Anti-Classical Swine Fever Virus Strategies. Microorganisms, 2021, 9(4): 761.

4. Zou X, Yang Y, Lin F, et al. Lactate facilitates classical swine fever virus replication by enhancing cholesterol biosynthesis. iScience, 2022, 25(11): 105353.

5. Bai J S, Zou L K, Liu Y Y, et al. Classical swine fever virus utilizes stearoyl-CoA desaturase 1-mediated lipid metabolism to facilitate viral replication. Journal of Virology, 2025, 99(6): e0055125.

6. Zhou B, Classical Swine Fever in China-An Update Minireview. Frontiers in Veterinary Science 2019, 6:8.

7. Bohara V S, Kumar S. Decoding the metabolic crosstalk between glycolysis and RNA viral pathogenesis. Virology, 2026, 615: 110766.

8. Chen S S, Qin T X, Luo S R, et al. Lactylation and viral infections: A novel link between metabolic reprogramming and immune regulation. PLoS Pathogens, 2025, 21(7): e1013366.

9. Zheng X R, Wang Y D, Zhang L H, et al. Construction of Macrophage Cell Lines Stably Expressing CSFV NS3-NS4A and NS3pro-NS4A Proteins and Proteomic Analysis. Chinese Journal of Animal and Veterinary Sciences, 2025, 56(11): 5839–5851.

10. Yang B, Hao Y, Yang J, et al. PI3K-Akt pathway-independent PIK3AP1 identified as a replication inhibitor of the African swine fever virus based on iTRAQ proteomic analysis[J]. Virus Research, 2023, 327: 199052.

11. Blanco J, Cameirao C, López M C, et al. Phosphatidylinositol-3-kinase-Akt pathway in negative-stranded RNA virus infection: a minireview. Archives of Virology, 2020, 165(10): 2165–2176.

12. Sun J, Yu H, Wang Y, et al. Classical swine fever virus NS5A protein activates autophagy via the PP2A-DAPK3-Beclin 1 axis. Journal of Virology, 2023, 97(12): e0098823.

13. Li J, Chen Z, Jin M, et al. Histone H4K12 lactylation promotes malignancy progression in triple-negative breast cancer through SLFN5 downregulation. Cellular Signalling, 2024, 124: 111468.

14. Wei E, Ji D, Jia Y, et al. BRD9 recognizes lactate-induced H3K18 lactylation to drive oncogenic chromatin remodeling in hepatocellular carcinoma[J]. Cell Death and Differentiation, 2026.

15. He W, Huang X, Zhang R, et al. Metabolic regulation of gene expression by histone lactylation. Nature, 2019, 574(7779): 575-580.

16. Zhang Y, Zhang H, Zhang X, et al. H3K18 lactylation marks tissue-specific active enhancers[J]. Genome Biology, 2022b, 23(1): 228.

17. Wang R, Wu Y, Zhou Z, et al. Benchmark of chromatin-protein interaction methods in haploid round spermatids. Frontiers in Cell and Developmental Biology, 2025, 13: 1572405.

18. Kaya-Okur H S, Wu S J, Codomo C A, et al. CUT&Tag for efficient epigenomic profiling of small samples and single cells. Nature Communications, 2019, 10(1): 1930.

19. Fu Z, Jiang S, Sun Y, et al. Cut&tag: a powerful epigenetic tool for chromatin profiling. Epigenetics, 2024, 19(1): 2293411.

20. Zhang S X, Ge L X, Wu S H, et al. Construction and Preliminary Analysis of CUT&Tag Library from Cambial Cells of Hevea brasiliensis Induced by Coronatine for Secondary Laticifer Differentiation. Chinese Journal of Tropical Crops, 2024, 45(10): 2010–2024.

21. Li F, Si W, Xia L, et al. Positive feedback regulation between glycolysis and histone lactylation drives oncogenesis in pancreatic ductal adenocarcinoma. Molecular Cancer, 2024, 23(1): 90.

22. Gockel J, Ramón-Zamorano G, Kimmel J, et al. CUT&Tag and DiBioCUT&Tag enable investigation of the AT-rich epigenome of Plasmodium falciparum from low-input samples. Cell Reports Methods, 2025, 5(8): 101110.

23. Zhu W H, Zeng S, Zhu S Q, et al. Histone H2B lysine lactylation modulates the NF-κB response via KPNA2 during CSFV infection[J]. International Journal of Biological Macromolecules, 2025, 299: 139973.

